# A Pubic Hair Is 172 Times More Pubic Than a Scalp Hair

**DOI:** 10.64898/2026.06.25.734686

**Authors:** Norichika Ogata, Tomoko Matsuda

## Abstract

Human hair is a common contaminant in GMP-controlled manufacturing environments, and its identification is important for contamination source investigation and corrective action. Because human hair can originate from multiple body sites, it is often necessary to determine not only the species of origin but also the anatomical source of the hair. Conventional forensic approaches distinguish scalp hair from body hair by microscopic examination of cuticle patterns, medullary structure, cross-sectional morphology, and pigment distribution. However, these methods depend on examiner expertise, are difficult to apply to damaged specimens, and provide limited quantitative information. In this study, we developed a proteomics-based approach for distinguishing scalp hair from pubic hair using identical sample preparation and analytical workflows. Comparative proteomic analysis identified keratin-associated proteins KAP 4-3 and KAP 9-6 as enriched in scalp hair, whereas cuticular keratins Ha7 and Ha8 were strongly enriched in pubic hair. Amino acid composition analysis further revealed that scalp hair-enriched proteins were highly cysteine-rich, consistent with sulfur-rich cross-linking matrix proteins, whereas pubic hair-enriched proteins exhibited characteristics of structural keratin filaments. These results demonstrate that proteomic signatures can provide a quantitative and objective means of determining the anatomical origin of human hair and may contribute to contamination source tracing in GMP manufacturing and forensic investigations.

## Introduction

Recent advances in mass spectrometry-based proteomics have enabled quantitative analyses of biological materials that were previously difficult to characterize. In biopharmaceutical manufacturing, host cell proteins (HCPs) are recognized as critical process-related impurities. Historically, HCP analysis focused primarily on the total burden of residual proteins, whereas modern proteomic approaches allow the identification and quantification of thousands of individual proteins. Such advances have significantly improved the characterization and control of biopharmaceutical products [1].

In addition to final product testing, Good Manufacturing Practice (GMP) requires comprehensive control of manufacturing environments and processes. Investigation of foreign particulate contamination is therefore an important component of pharmaceutical quality systems [2]. Among the various contaminants encountered in manufacturing environments, human hair is a well-recognized source of contamination [3]. Identification of the anatomical origin of contaminating hair may facilitate root-cause analysis because different hair types are associated with different contamination pathways. For example, contamination by scalp hair may indicate inadequate hair-covering procedures, whereas contamination by pubic hair may suggest deficiencies in gowning practices, changing-room procedures, or personal hygiene management. Determining the anatomical source of human hair contaminants may therefore contribute to the prevention of recurrent contamination events.

Conventional approaches for identifying the origin of human hair have largely been developed within forensic science. These methods typically rely on microscopic examination of cuticle patterns, medullary structures, cross-sectional morphology, and pigment distribution. Although effective in many cases, such approaches are dependent on examiner expertise, may be difficult to apply to damaged specimens, and provide limited quantitative information [4–8].

Previous studies have demonstrated that hair fibers are composed primarily of low-sulfur α-keratins and sulfur-rich matrix proteins. Keratin-associated proteins (KAPs), which constitute major sulfur-rich matrix proteins, are known to fill the spaces between keratin filaments and contribute to mechanical strength and shape retention through extensive disulfide cross-linking. In contrast, specific cuticular keratins, including hHa7, have been reported to be highly expressed in sexual hairs [9–13]. These findings suggest that scalp hair and pubic hair possess distinct protein compositions and may therefore be distinguishable by proteomic analysis.

Comprehensive characterization of hair proteomes remains challenging because extensive cross-linking and the highly insoluble nature of hair proteins limit extraction efficiency [14,15]. However, complete proteome coverage may not be necessary for practical discrimination of hair origin. Relative differences detectable under a standardized analytical workflow may be sufficient to classify hair according to its anatomical source.

In the present study, we investigated whether SWATH-MS–based proteomic profiling could distinguish scalp hair from pubic hair using a common sample preparation workflow. By comparing proteins identified from both hair types, we sought to evaluate the feasibility of proteomics-based hair origin determination and to identify candidate protein markers applicable to contamination source tracing in GMP-regulated manufacturing environments.

## Methods

### Protein extraction from hair samples

Approximately 0.04 g of scalp hair and pubic hair were independently processed using the same extraction protocol. Hair samples were sequentially washed with Milli-Q water, 99.5% ethanol, and acetone in 50-mL tubes to remove surface contaminants. The washed samples were air-dried on Kimwipes and cut into small fragments using dissecting scissors. Fragmented hair samples were transferred to PROKEEP protein low binding tubes (WATSON, PK-15C-500N). For protein extraction, 7.7 mg of dithiothreitol (DTT) was added using Pierce™ DTT (Dithiothreitol), No-Weigh™ Format (Thermo Fisher Scientific, A39255). DTT was dissolved in 6.4 M guanidine hydrochloride containing 50 mM Tris-HCl, and additional extraction buffer was added to obtain a final volume of 1 mL. Samples were incubated at 60°C for 6 h in a heat block, with vortex mixing performed once during incubation. After incubation, samples were centrifuged at 16,000 × g for 10 min. Buffer exchange was performed using Zeba™ Spin Desalting Columns, 7K MWCO (Thermo Fisher Scientific, 89883). Columns were equilibrated with 50 mM Tris-HCl (pH 7.9) by three successive washes with 300 μL buffer and centrifugation at 1,500 × g for 1 min. Subsequently, 130 μL of the extracted protein solution was loaded onto the column and centrifuged at 1,500 × g for 1 min. Because the eluate appeared turbid, an additional clarification step was performed by centrifugation at 16,000 × g for 5 min. The resulting supernatants were transferred to Protein LoBind tubes and stored at −80°C until further analysis.

### Protein quantification and enzymatic digestion

Protein concentrations were determined using Pierce™ Dilution-Free™ Rapid Gold BCA Protein Assay (Thermo Fisher Scientific, A55860). The measured protein concentrations were 3.3 mg/mL for the scalp hair extract and 1.5 mg/mL for the pubic hair extract. Protein digestion was performed using the EasyPep™ 96 Micro MS Sample Prep Kit (Thermo Fisher Scientific, A57864) according to the manufacturer’s protocol. Samples were adjusted to a final volume of 10 μL using the supplied lysis buffer. Specifically, 3.0 μL of scalp hair extract and 6.7 μL of pubic hair extract were mixed with lysis buffer to a final volume of 10 μL. Subsequently, 5 μL of Reduction Solution and 5 μL of Alkylation Solution were added sequentially. Samples were incubated at 95°C for 5 min. Trypsin supplied with the kit (5 μL) was then added, and samples were incubated at 37°C for 3 h with shaking at 800 rpm. Digestion was terminated by addition of 5 μL of Digestion Stop Solution. Digested peptide samples were stored at −80°C until cleanup.

### Peptide cleanup

Peptide cleanup was performed using the EasyPep™ 96 Micro MS Sample Prep Kit (Thermo Fisher Scientific, A57864). Digested samples were applied to the cleanup columns and centrifuged at 1,000 rpm for 3 min. Columns were washed once with 100 μL Solution A and three times with 100 μL Solution B, with centrifugation at 2,000 rpm for 2 min after each wash step. Peptides were eluted with 100 μL Elution Solution by centrifugation at 2,000 rpm for 2 min. Eluates were transferred to Protein LoBind tubes and dried using an evaporator at 37°C for 1 h. Dried peptide samples were stored at −80°C until LC–MS/MS analysis.

### LC–MS/MS analysis

Prior to analysis, dried peptide samples were reconstituted in 25 μL of 0.1% formic acid. The peptide solutions were filtered and subjected to LC–MS/MS analysis. Data-independent acquisition (SWATH-MS) was performed for comparative proteomic analysis of scalp hair and pubic hair samples. We analyzed proteome data as previously described [1]. Referential protein sequences were obtained from NCBI’s RefSeq Genomes [16]. A modified MA plot was generated using log10(Pubic) + log10(Scalp) as the abundance axis and log10(Pubic) − log10(Scalp) as the differential abundance axis.

## Results and Discussions

A total of 321 proteins were identified in this study (Table 1). Among the identified proteins, keratin, type I cuticular Ha7 (NP_003761.3) and keratin, type I cuticular Ha8 (NP_006762.3) were markedly enriched in pubic hair, exhibiting 172-fold and 107-fold higher abundances, respectively, than in scalp hair. Conversely, keratin-associated protein (KAP) 9-6 (NP_001264260.1) and keratin-associated protein (KAP) 4-3 (NP_149443.1) were enriched in scalp hair, showing approximately 8-fold higher abundances than in pubic hair (Figure 1). These findings indicate substantial differences in protein composition between scalp hair and pubic hair.

**Table 1.**
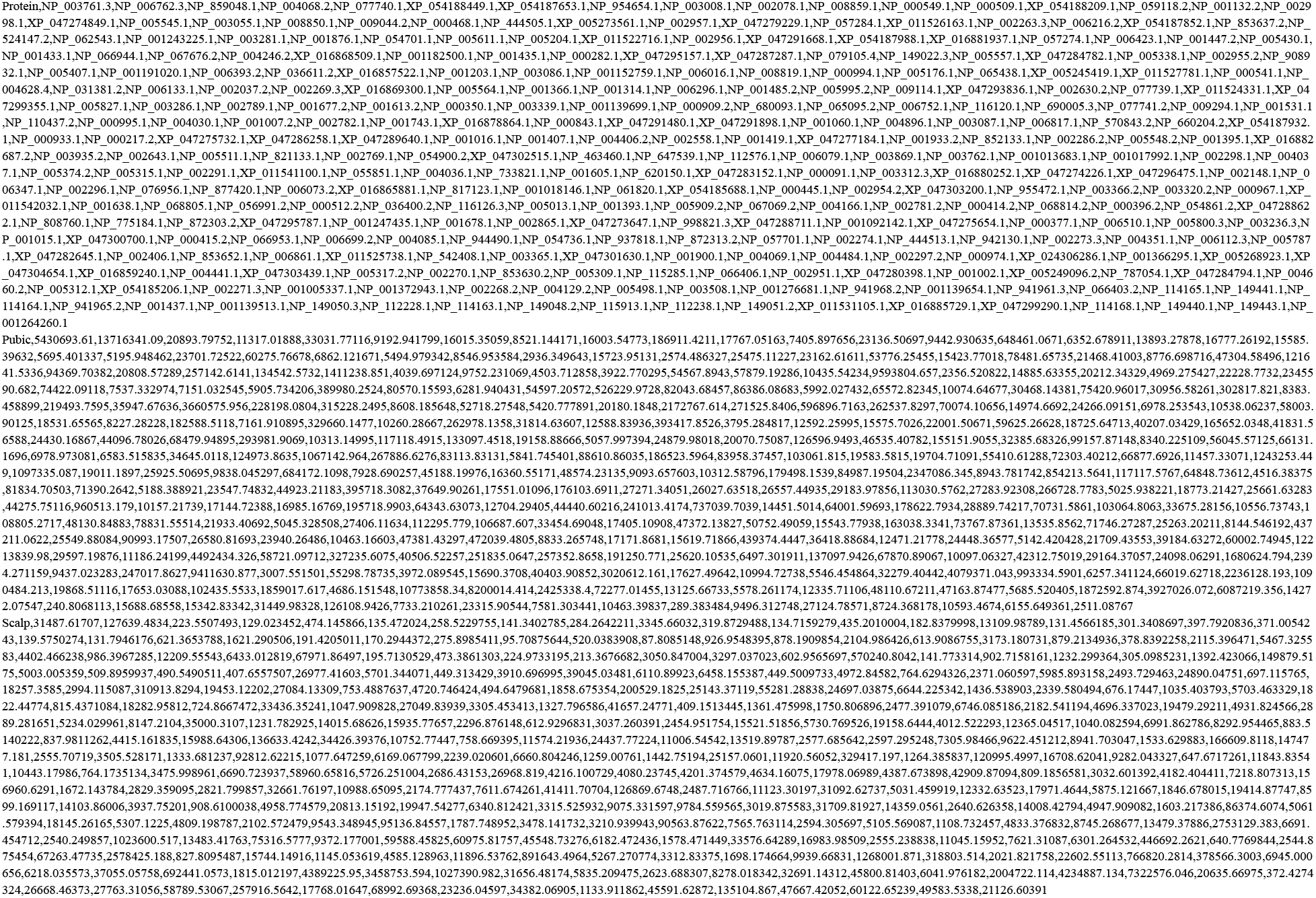
SWATH-MS peak areas of proteins detected in scalp hair and pubic hair.

**Figure 1.**
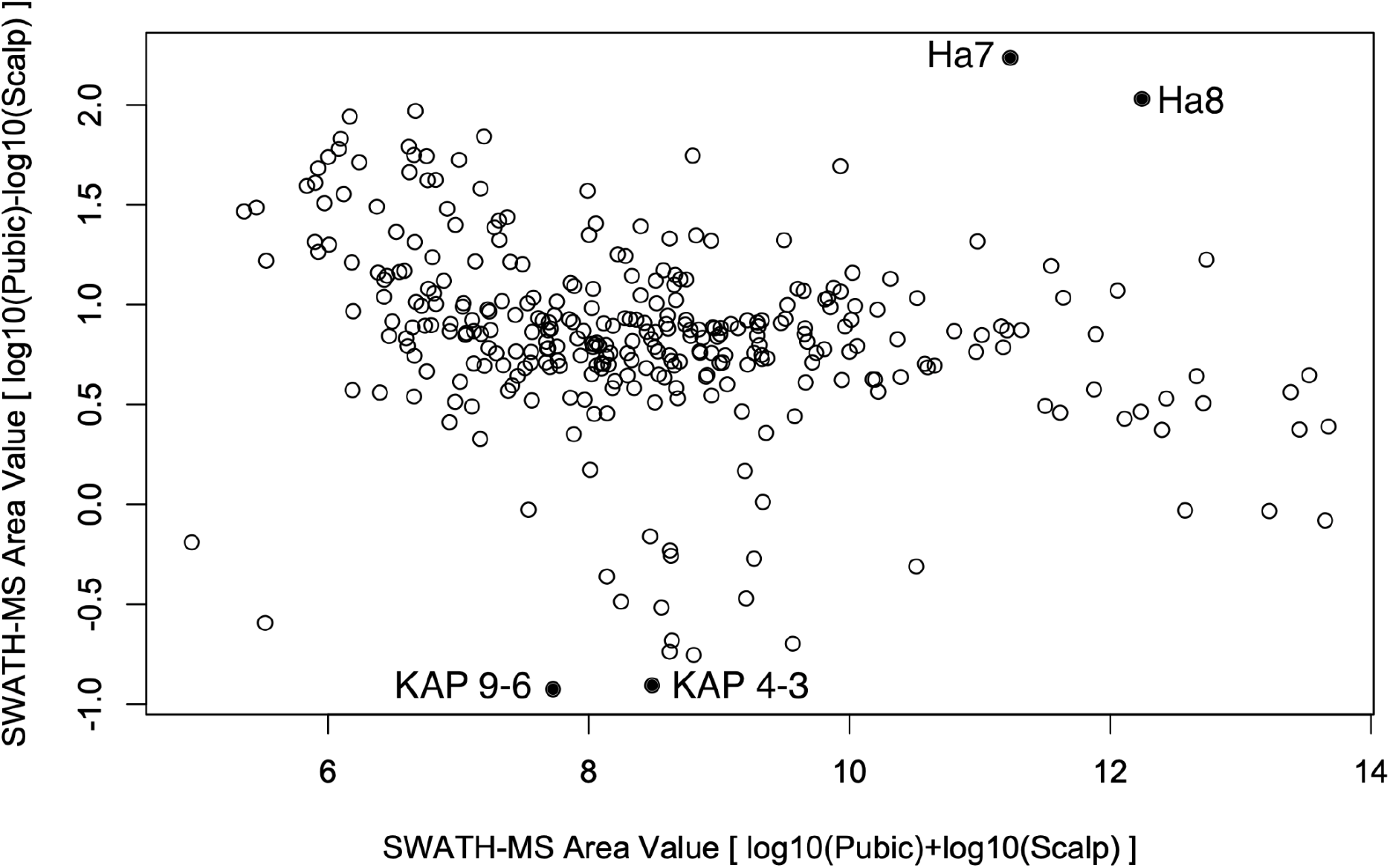
Comparative MA plot of proteins identified in scalp hair and pubic hair samples by SWATH-MS Each point represents a single protein identified in both samples. The x-axis indicates the overall protein abundance calculated as log10(Pubic) + log10(Scalp), while the y-axis represents the relative abundance difference calculated as log10(Pubic) − log10(Scalp). Positive y-values indicate proteins enriched in pubic hair, whereas negative y-values indicate proteins enriched in scalp hair. Keratin, type I cuticular Ha7 (NP_003761.3) and Ha8 (NP_006762.3) were among the proteins most strongly enriched in pubic hair, whereas keratin-associated protein 4-3 (NP_149443.1) and keratin-associated protein 9-6 (NP_001264260.1) were among the proteins most strongly enriched in scalp hair. These proteins are indicated by filled black circles.

The proteins enriched in scalp hair and pubic hair belonged to distinct structural classes. Scalp hair was characterized by increased abundance of keratin-associated proteins, whereas pubic hair was enriched in cuticular keratins. These findings suggest that the two hair types differ not only quantitatively but also in their underlying structural protein architecture.

Amino acid compositions were calculated from RefSeq protein sequences. Amino acid sequence analysis revealed substantial compositional differences between proteins enriched in scalp hair and those enriched in pubic hair. KAP4-3 and KAP9-6 contained exceptionally high proportions of cysteine residues (approximately 31–36%), whereas Ha7 and Ha8 contained only 5–6% cysteine. The cysteine-rich nature of scalp hair-associated proteins is consistent with their proposed role as sulfur-rich matrix proteins involved in disulfide-mediated cross-linking and mechanical stabilization of the hair shaft.

Previous studies have reported that keratin-associated proteins constitute major sulfur-rich matrix components of hair fibers and contribute to filament cross-linking through disulfide bond formation. In contrast, hHa7 has been reported to be preferentially expressed in sexual hairs. The present observations are therefore consistent with previously reported biological differences between scalp hair and sexual hairs and provide direct proteomic evidence supporting these distinctions.

Comprehensive characterization of hair proteomes remains challenging because extensive disulfide cross-linking and protein insolubility limit extraction efficiency. Despite these limitations, clear discrimination between scalp hair and pubic hair was achieved in the present study. This finding suggests that complete proteome coverage may not be necessary for practical determination of hair origin.

From a practical perspective, the present approach may contribute to contamination source tracing in GMP-regulated manufacturing environments. Different hair types may indicate different contamination pathways, and determination of hair origin could facilitate root-cause investigations and implementation of corrective and preventive actions (CAPA). Unlike conventional forensic approaches based primarily on microscopic observation, proteomic profiling provides quantitative and objective molecular evidence that may complement existing methods.

Taken together, the results demonstrate that scalp hair and pubic hair possess distinct proteomic signatures. In particular, Ha7 and Ha8 represent candidate markers of pubic hair, whereas KAP4-3 and KAP9-6 represent candidate markers of scalp hair. These proteins may serve as practical indicators for anatomical origin determination and contamination source tracing. Future studies using multiple donors will be necessary to evaluate inter-individual variation.

## Conclusion

SWATH-MS analysis identified distinct proteomic signatures in scalp hair and pubic hair. Ha7 and Ha8 were highly enriched in pubic hair, whereas KAP4-3 and KAP9-6 were enriched in scalp hair. These findings demonstrate the feasibility of proteomics-based determination of hair origin and suggest potential applications in GMP contamination source tracing.

## References

1. Ogata N, Matsuda T, Hosaka A, Shina A, Hashiba N, Uchida K, et al. Database chemistry for genomics-based safety and quality evaluation of biologics. bioRxiv. bioRxiv; 2025. doi:10.1101/2025.02.14.638017

2. Active Pharmaceutical Ingredients Committee. Guidance on handling of insoluble matter and foreign particles in APIs. Brussels: CEFIC; 2015. Available: https://www.gmp-navigator.com/mygmp/wirk-und-hilfsstoffe/guidelines-active-pharmaceutical-ingredients?file=files/eca/userFiles/mygmp-guidelines/20150626foreignparticleguideline_final.PDF

3. Hair As A Pharmaceutical Contaminant: How It Is Identified, And How We Can Tell Where It Originated - Analysis Solutions. In: Gateway Analytical [Internet]. 5 Jan 2023 [cited 25 Jun 2026]. Available: https://gatewayanalytical.com/hair-as-a-pharmaceutical-contaminant/?utm_source=chatgpt.com

4. [Active Pharmaceutical Ingredients Committee]. Guidance on Handling of Insoluble Matter and Foreign Particles in APIs.

5. Beckert JC. Forensic hair microscopy. Handbook of Trace Evidence Analysis. Wiley; 2020. pp. 219–321. doi:10.1002/9781119373438.ch4

6. Forensic Hair Comparison: Background Information for Interpretation. [cited 25 Jun 2026]. Available: https://www.ojp.gov/ncjrs/virtual-library/abstracts/forensic-hair-comparison-background-information-interpretation

7. D’Orio E, ResearchCentre BF, Calabrese G, Lucanto C, Montagna P, ResearchCentre BF, et al. Assessing performance in forensic hair examination: A review. International Journal of Law in Changing World. 2023;2: 102–119.

8. Ryu SR, Suh J, Kim H, Shin K. The surface and internal features of pubic hair: A comparative study with those of scalp hair. Exp Dermatol. 2023;32: 1509–1520.

9. Shimomura Y, Ito M. Human hair keratin-associated proteins. J Investig Dermatol Symp Proc. 2005;10: 230–233.

10. Parry DAD, Smith TA, Rogers MA, Schweizer J. Human hair keratin-associated proteins: sequence regularities and structural implications. J Struct Biol. 2006;155: 361–369.

11. Rogers MA, Langbein L, Winter H, Beckmann I, Praetzel S, Schweizer J. Hair keratin associated proteins: characterization of a second high sulfur KAP gene domain on human chromosome 21. J Invest Dermatol. 2004;122: 147–158.

12. Jave-Suarez LF, Langbein L, Winter H, Praetzel S, Rogers MA, Schweizer J. Androgen regulation of the human hair follicle: the type I hair keratin hHa7 is a direct target gene in trichocytes. J Invest Dermatol. 2004;122: 555–564.

13. Rouse JG, Van Dyke ME. A review of keratin-based biomaterials for biomedical applications. Materials (Basel). 2010;3: 999–1014.

14. Lee YJ, Rice RH, Lee YM. Proteome analysis of human hair shaft: from protein identification to posttranslational modification. Mol Cell Proteomics. 2006;5: 789–800.

15. Adav SS, Subbaiaih RS, Kerk SK, Lee AY, Lai HY, Ng KW, et al. Studies on the proteome of human hair - identification of histones and deamidated keratins. Sci Rep. 2018;8: 1599.

16. NCBI’s RefSeq Genomes [cited 25 Jun 2026]. Available: https://ftp.ncbi.nlm.nih.gov/genomes/all/GCF/000/001/405/GCF_000001405.26_GRCh38/GCF_000001405.26_GRCh38_protein.faa.gz

